# Open multimodal iEEG-fMRI dataset from naturalistic stimulation with a short audiovisual film

**DOI:** 10.1101/2021.06.09.447733

**Authors:** Julia Berezutskaya, Mariska J. Vansteensel, Erik J. Aarnoutse, Zachary V. Freudenburg, Giovanni Piantoni, Mariana P. Branco, Nick F. Ramsey

## Abstract

Intracranial human recordings are a valuable and rare resource that the whole neuroscience community can benefit from. Making such data available to the neuroscience community not only helps tackle the reproducibility issues in science, it also helps make more use of this valuable data. The latter is especially true for data collected using naturalistic tasks. Here, we describe a dataset collected from a large group of human subjects while they watched a short audiovisual film. The dataset is characterized by several unique features. First, it combines a large amount of intracranial data from 51 intracranial electroencephalography (iEEG) participants, who all did the same task. Second, the intracranial data are accompanied by fMRI recordings acquired for the same task in 30 functional magnetic resonance imaging (fMRI) participants. Third, the data were acquired using a rich audiovisual stimulus, for which we provide detailed speech and video annotations. This multimodal dataset can be used to address questions about neural mechanisms of multimodal perception and language comprehension as well as the nature of the neural signal acquired during the same task across brain recording modalities.

## Background & Summary

We live in the world of data, and big high-quality datasets that lend themselves to modern sophisticated analyses are becoming increasingly sought-after. Following examples in other branches of science, cognitive neuroscience is adopting an ever-growing trend for open science and data sharing^1–3^. Lots of datasets from volunteers participating in cognitive neuroscience experiments are now becoming publicly available^4–9^. This is coupled with a recent trend to use more naturalistic designs in these experiments as they provide rich versatile datasets and as such lend themselves well to application of many different analyses targeting various aspects of complex cognition^10–12^. The open research practices promote data reuse, research reproducibility, scientific collaboration and novel ways to analyze the data that have not been possible before^3, 13^ and therefore have the potential to advance the entire field of cognitive neuroscience forward faster than ever before.

Cognitive neuroscience experiments are concerned with the neural mechanisms of cognitive processes including speech, sensory perception, memory, social interactions and others. These are most often studied with popular techniques, such as functional magnetic resonance imaging (fMRI), electroencephalography (EEG) and magnetoencephalography (MEG) and current openly available datasets indeed contain data collected with these techniques^4–9, 14, 15^. Despite their great value for the field, these non-invasive techniques have a number of important limitations, such as lack of temporal resolution in fMRI^16, 17^, lack of spatial resolution in EEG and MEG and susceptibility to artefacts that render part of the recorded signal unusable (EEG and MEG)^18–20^. To study the highly dynamic cognitive processes in humans (speech, in particular), techniques that provide high spatial and temporal resolution, and clean neural signal are preferred.

One such technique is human intracranial electroencephalography (iEEG). IEEG data are collected from patients who participate in a relatively rare procedure for localization of the source of their epileptic seizures. For this, patients are implanted subdurally with electrode grids (electrocorticography, ECoG) and/or depth electrodes (stereo-electroencephalography, sEEG) typically for a week of clinical monitoring, during which the patients can also participate in research experiments. Direct contact with the brain tissue grants iEEG several advantages compared to non-invasive brain recording modalities, including a combination of high temporal and spatial resolution and exceptional signal-to-noise ratio. Human iEEG research has contributed to our fundamental understanding of high-level cognition that cannot be studied in animals, such as speech^21–24^, semantic and concept representation^25–27^ and abstract thought^28^. In addition, iEEG research on speech has shown significant promise for the development of advanced brain-computer interfaces aimed to restore communication in paralyzed patients^29–32^.

The unique characteristics of human iEEG recordings make them a a valuable resource of information about the brain that should be used to the most of its potential through data sharing and open collaborations. However, due to multiple factors, the data are rarely shared. First, iEEG can only be obtained in the clinical setting, and as such, it is difficult and slow to collect for research purposes. Few medical centers in the world that acquire iEEG data suffer from low patient rates (5-10 a year) that, together with variability in electrode coverage, cause long study timeframes and low sample sizes. Moreover, iEEG is sensitive medical data and many centers lack ethical protocols that allow public sharing. As a result, there has been little publicly available iEEG data so far (with a few notable exceptions^33–35^).

Our lab has been collecting iEEG data for over ten years. A few years ago we developed ethical protocols that addressed the issue of data sharing and allowed us to request patients’ consent to publish their de-identified data and allow open access thereof to the entire research community. The possibility of data sharing was further facilitated by progress our colleagues made on a new standard iEEG data format - iBIDS^36^ that greatly simplifies and unifies data curation and preparation for public sharing.

As a result of this work, we here present the first large multimodal iEEG-fMRI dataset from a naturalistic cognitive task. The present dataset is unique in a number of ways. First, it contains a large amount of iEEG data (51 subjects who all did the same task). Second, the dataset provides additional fMRI recordings (30 subjects) from the same task. Eighteen subjects did the task with both iEEG and fMRI. Third, the data comes from naturalistic stimulation with a short audiovisual film, for which we provide rich audio and video annotations. Inclusion of data from two neural recording modalities opens up new possibilities for research on neurovascular coupling in a context of a naturalistic experiment.

The dataset we present can be used to target many theoretical, methodological and applied questions in cognitive neuroscience. We believe this work has a potential to promote open science and sharing in the iEEG field and support open research practices in the cognitive neuroscience community as a whole.

## Methods

### Participants

All participants were admitted to the University Medical Center Utrecht for diagnostic procedures related to their medication-resistant epilepsy. They underwent intracranial electrode implantation to determine the source of seizures and test the possibility of surgical removal of the corresponding brain tissue. The tasks (movie watching and resting state) were performed by the patients as part of clinical function mapping procedures, in which our team was involved, or as part of their participation in scientific research done by our group. In the latter case, patients gave their written informed consent to participate in such research. All patients gave a written permission to use their data for research purposes and to share their de-identified data publicly. For participants under 18, the informed consent was obtained from the participant’s parents and/or legal guardian. If older than 12, these participants also signed the informed consent form. The study was approved by the Medical Ethical Committee of the Utrecht University Medical Center in accordance with the Declaration of Helsinki (2013).

#### IEEG participants

Data from fifty-one iEEG patients (age 25±15, 32 females) are included in the present dataset. Basic demographic information about all participants in the dataset is shown in Table 1. Forty-six patients were implanted with subdural ECoG grids (clinical grids with 2.3 mm exposed diameter, inter-electrode distance 10 mm, between 48 and 128 contact points). Six patients were additionally implanted with a high-density ECoG grid (with 1.3 mm exposed diameter, inter-electrode distance 3-4 mm, with 32, 64 or 128 contact points). Sixteen patients were implanted with sEEG electrodes (between 4 and 173 contact points). Most patients had perisylvian grid coverage and most had electrodes in frontal and motor cortices (Figure 1c).

**Table 1.** Demographic information about all participants in the dataset (M: male, F: female, R: right, L: left). Column ’Language’ refers to the language-dominant hemisphere, determined using one of the standard techniques (column ’Language Technique’). The following techniques were used: fMRI (functional magnetic resonance imaging), fTCD (functional transcranial Doppler sonography), Wada (intracarotid amobarbital test), ECS (electrocortical stimulation). Entry ’n/a’ indicates missing information. For each participant, columns ’IEEG’ and ’FMRI’ indicate whether data of the corresponding modality is present in the dataset. Column ’IEEG Hemisphere’ indicates the hemisphere, in which iEEG electrodes were implanted for patients with iEEG recordings available. Column ’HD’ indicates availability of high-density iEEG recordings.

| ID | Sex | Age | Handedness | Language | Language Technique | IEEG | FMRI | IEEG Hemi-sphere | HD |
| --- | --- | --- | --- | --- | --- | --- | --- | --- | --- |
| sub-01 | M | 55 | R | n/a | n/a | yes | no | LR | no |
| sub-02 | F | 9 | R | L | fTCD + fMRI | yes | no | L | no |
| sub-03 | F | 33 | R | L | Wada | yes | no | L | no |
| sub-04 | M | 11 | R | possibly L | fMRI | no | yes | n/a | n/a |
| sub-05 | F | 33 | R | L | fTCD | yes | no | L | no |
| sub-06 | F | 43 | R | L | fMRI | yes | no | L | no |
| sub-07 | F | 42 | R | L | fMRI | yes | yes | L | no |
| sub-08 | M | 17 | R | L | fTCD + fMRI | no | yes | n/a | n/a |
| sub-09 | F | 33 | R | L | fTCD | yes | yes | L | no |
| sub-10 | M | 8 | R | L | Wada | yes | no | L | no |
| sub-11 | F | 9 | n/a | n/a | n/a | no | yes | n/a | n/a |
| sub-12 | M | 37 | R | L | ECS | yes | no | L | no |
| sub-13 | F | 17 | L | R | Wada | yes | yes | R | no |
| sub-14 | F | 18 | R | R | Wada | yes | yes | L | yes |
| sub-15 | M | 14 | R | L | fMRI | no | yes | n/a | n/a |
| sub-16 | M | 17 | R | L | Wada | yes | yes | L | no |
| sub-17 | M | 28 | L | L | Wada | yes | no | R | no |
| sub-18 | F | 15 | R | L | fMRI | yes | yes | L | no |
| sub-19 | F | 8 | R | possibly L | fMRI | yes | no | L | no |
| sub-20 | F | 25 | L | L | Wada | yes | no | L | no |
| sub-21 | F | 51 | R | L | Wada | yes | no | R | no |
| sub-22 | M | 21 | R | L | fMRI | yes | yes | L | no |
| sub-23 | M | 40 | R | inconclusive | fTCD | yes | no | LR | no |
| sub-24 | F | 47 | R | L | Wada | yes | yes | L | no |
| sub-25 | M | 14 | R | L | fTCD | yes | no | L | no |
| sub-26 | F | 48 | R | L | fMRI | yes | no | L | no |
| sub-27 | M | 15 | R | L | fMRI | yes | yes | L | no |
| sub-28 | M | 21 | R | L | fMRI | yes | yes | R | no |
| sub-29 | F | 23 | L | possibly bilat-<br>eral | fTCD | no | yes | n/a | n/a |
| sub-30 | M | 24 | R | n/a | n/a | yes | no | L | yes |
| sub-31 | F | 13 | L | R | Wada | yes | yes | LR | no |
| sub-32 | F | 6 | R | L | Wada | yes | no | L | no |
| sub-33 | F | 9 | R | n/a | n/a | yes | no | R | no |
| sub-34 | F | 51 | R | L | ECS | yes | no | L | no |
| sub-35 | F | 14 | R | possibly L | fMRI | no | yes | n/a | n/a |
| sub-36 | F | 52 | R | L | fMRI | yes | no | L | yes |
| sub-37 | M | 5 | n/a | L | Wada | yes | no | L | no |
| sub-38 | F | 14 | R | R | fMRI | yes | no | L | no |
| sub-39 | F | 5 | R | possibly L | ECS (fTCD<br>inconclu-<br>sive, maybe<br>bilateral) | yes | no | L | no |
| sub-40 | M | 49 | R | L | fMRI | yes | no | L | no |
| sub-41 | M | 7 | originally R,<br>but switched<br>to L | possibly L | ECS (fMRI in-<br>conclusive) | yes | yes | L | no |
| sub-42 | F | 23 | R | L | fMRI (fTCD<br>inconclusive) | yes | no | R | no |
| sub-43 | M | 19 | R | L | Wada | yes | yes | R | yes |
| sub-44 | M | 26 | R | L | Wada | no | yes | n/a | n/a |
| sub-45 | M | 19 | R | L | fMRI | yes | yes | L | yes |
| sub-46 | F | 41 | L | L | fMRI | yes | yes | L | no |
| sub-47 | F | 22 | L | R | fTCD + fMRI | no | yes | n/a | n/a |
| sub-48 | F | 18 | R | L | fTCD and/or<br>fMRI | yes | no | L | no |
| sub-49 | F | 10 | n/a | L | Wada | yes | no | L | no |
| sub-50 | F | 9 | R | bilateral | fMRI | yes | no | L | no |
| sub-51 | M | 46 | R | L | fMRI + fTCD | yes | yes | L | no |
| sub-52 | M | 17 | R | L | unspecified | no | yes | n/a | n/a |
| sub-53 | M | 14 | L | possibly L | fMRI | no | yes | n/a | n/a |
| sub-54 | F | 31 | R | L | fMRI | yes | no | L | no |
| sub-55 | F | 23 | R | L | Wada | yes | yes | L | no |
| sub-56 | M | 18 | R | L | fMRI | no | yes | n/a | n/a |
| sub-57 | F | 36 | R | L | fMRI | yes | no | L | no |
| sub-58 | F | 16 | R | L | fMRI | yes | no | L | no |
| sub-59 | F | 30 | L | L | fMRI | yes | no | L | no |
| sub-60 | M | 42 | R | L | Wada | yes | yes | L | yes |
| sub-61 | F | 16 | R | L | fTCD | yes | no | L | no |
| sub-62 | F | 14 | R | possibly L | fMRI | no | yes | n/a | n/a |
| sub-63 | M | 5 | n/a | L | ECS | yes | no | L | no |

**Figure 1.**
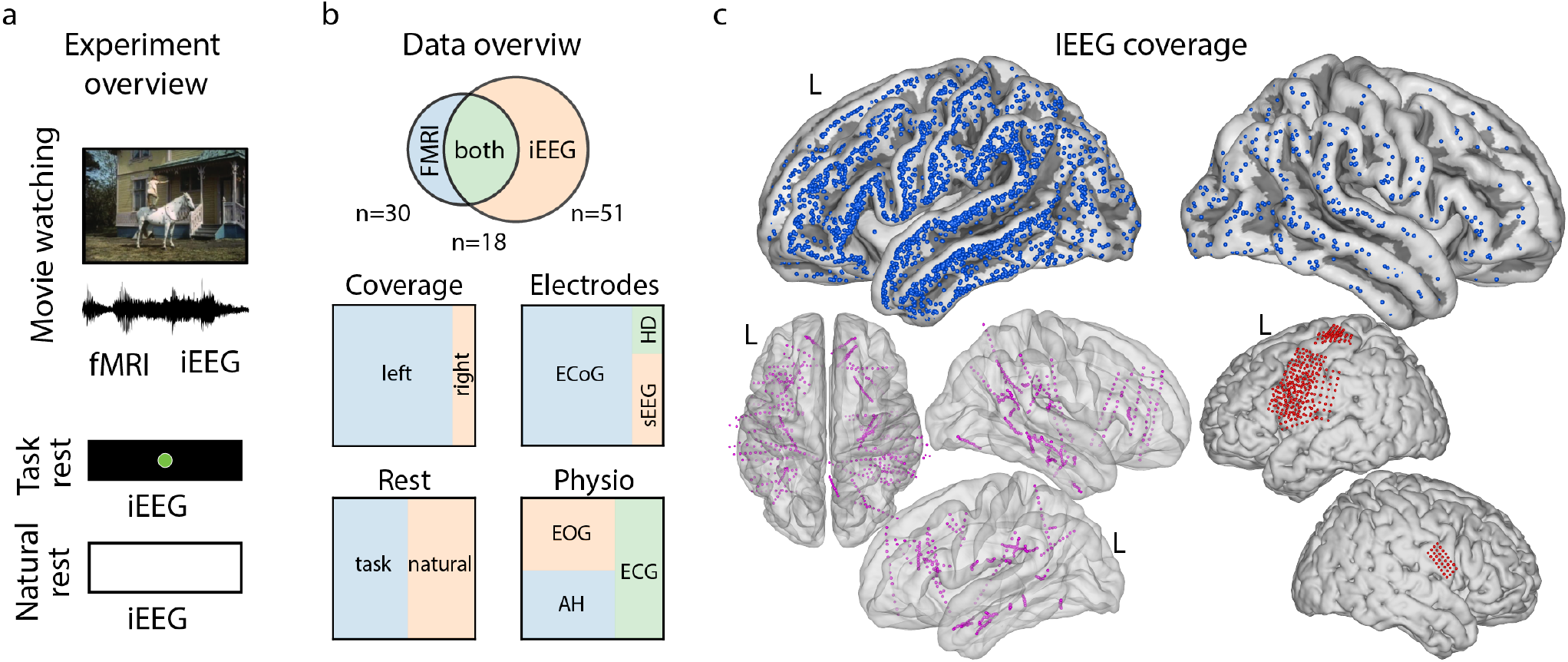
Overview of experiments and data. a. Overview of two experiments: movie-watching (iEEG and fMRI) and rest (iEEG). If rest task was not available, rest data was taken from natural rest in continuous 24/7 iEEG recordings. b. Data overview with number of subjects per brain recording modality (*n*). Tree partition plots below show iEEG data proportions in hemisphere coverage (left, right), electrode type (clinical ECoG, HD ECoG, sEEG), type of rest (task rest, natural rest) and type of physiological recordings available (EOG, breathing - AH, ECG). c. IEEG electrode coverage. Top plots show ECoG electrode locations (in cyan) on the average surface. Bottom left plots show sEEG electrodes (in black), bottom right plots show HD ECoG electrodes (in red).

Forty-five patients were implanted with electrodes in the left hemisphere, which was also language dominant in most cases, based on fMRI, intracarotid amobarbital test (Wada), electrical stimulation or functional transcranial Doppler sonography test (Table 1). Nine patients had electrodes in the right hemisphere, and three patients had electrodes in both hemispheres.

#### FMRI participants

As part of the presurgical workup eighteen of the above fifty-one iEEG patients underwent fMRI recordings and participated in the fMRI version of the movie-watching task. In addition, there were eleven more patients who only participated in the fMRI experiment. In total, thirty participants were included (age 22±11, 14 females). The diagram showing the overlap between the fMRI and iEEG participants is shown in Figure 1b.

### Experimental procedures

#### Movie-watching experiment (iEEG and fMRI)

The short movie-watching task was developed as part of the standard battery of clinical tasks for presurgical functional language mapping. Therefore, most patients performed this task based on the clinical request. Remaining patients were offered the task as part of research they has agreed to participate in. Each participant watched the short movie either during the iEEG or the fMRI experiment. Some participants watched the movie during both iEEG and fMRI recordings. The fMRI experiment always preceded the iEEG recordings as fMRI data collection was part of the presurgical workup that typically took place several weeks prior to electrode implantation. In the movie-watching experiment, each patient was asked to attend to the short movie made of fragments from one of the Pippi Longstocking movies (see details below). No fixation cross was displayed in the middle of the screen or elsewhere. Instead, the participants were free to watch the film in as a naturalistic setting as possible. In the case of the fMRI experiment, the video was delivered on a screen through a scanner mirror and the audio was delivered through earphones. In the case of the iEEG experiment, the video was delivered on a computer screen (21 inches in diagonal) placed directly in the patient’s room (at approximately one meter to the patient’s face), and the stereo sound was delivered through speakers with the volume level adjusted for each patient.

In both cases the movie was presented using the Presentation software (Version 18.0, Neurobehavioral Systems) and the sound was synchronized with the neural recordings.

#### Resting state experiment (iEEG)

During iEEG recordings twenty-six iEEG patients participated in a resting state experiment. Some patients performed the resting state task and the movie-watching task on the same day, others did the tasks on different days. The movie-watching task was performed first if it was part of the clinical testing of functional cortical mapping. The resting state experiment was collected for research purpose only.

No resting state recordings during fMRI experiments were available.

#### Natural resting state data (iEEG)

Even though there was generally more time to collect resting state data with iEEG patients, it was not always feasible due to many reasons. Twenty four patients did not participate in a separate resting state task. In order to provide some form of baseline neural activity for the remaining iEEG patients, we selected a 3-minute fragment of ‘natural rest’ from each of these patients’ continuous 24/7 clinical iEEG recordings. We used clinical audiovisual recordings of the room to make sure that during ’natural rest’ patients did not speak and were not spoken to, it was quiet in the room and the patient was resting with their eyes open.

### Movie stimulus

A 6.5 minute short movie, made of fragments from “Pippi on the Run” (Pårymmen med Pippi Långstrump, 1970) was edited together to form a coherent plot with limited task duration. As a clinical task designed for language mapping, the movie consisted of 13 interleaved blocks of speech and music, 30 seconds each (seven blocks of music, six blocks of speech). The movie was originally in Swedish but dubbed into Dutch. Due to copyright permissions we cannot share the movie stimulus itself in a public repository. However, we are allowed to distribute it upon request to the corresponding author for strictly non-profit research purposes related to the present neural dataset. At the same time as part of the dataset we provide detailed annotations of the audio and video content of the movie stimulus.

#### Audio annotations

Annotation of the movie soundtrack was done manually using Praat^37^ (http://www.praat.org/). Onsets and offsets of several language features were annotated including phonemes, syllables, words, clauses and sentences. We also marked onsets and offsets of individual verbs due to their central role in the sentence structure. In addition, we annotated onsets and offsets of words spoken by each story character: Pippi, Annika, Tommi, Mom, Dad and Konrad.

#### Characters, scenes and higher-level concepts

Video annotations^27^ were obtained using the commercial deep neural network Clarify General. The network received video frames one by one and returned 20 visual concept labels that were most likely present in the frame. The network was pretrained on a large dataset using a dictionary of 5,000 unique visual concepts. The output of the visual concept recognition model was then manually corrected by removing irrelevant labels and adjusting incorrect assignments. The final list of labels consisted of 129 unique visual concepts. Most labels referred to objects present in the frame, for example, ‘house’, ‘table’, ‘animal’, ‘rock’, etc., some labels described a state or relation (such as ‘seated’, ‘equestrian’, ‘wooden’, ‘together’, ‘outdoors’, etc.) or action (such as ‘walk’, ‘travel’, ‘dance’, ‘climb’, ‘smile’, etc.).

In addition, we manually annotated presence of each story character in each frame.

### Data acquisition details

#### IEEG data acquisition

During the experiments, iEEG data were acquired with a 128 channel recording system (Micromed, Treviso, Italy). In most cases data were sampled at the rate of 512 Hz and filtered at 0.15–134.4 Hz (38 patients). In some cases data were sampled at the rate of 2048 Hz and filtered at 0.3–500 Hz (13 patients). An external reference electrode typically placed on the mastoid part of the temporal bone was used as signal reference. In addition to the clinical iEEG recordings, six paricipants were implanted with HD ECoG grids. In two HD participants, HD ECoG data was recorded at 2000 Hz (filtered at 0.3–500 Hz) with a Blackrock system (Blackrock Microsystems, https://www.blackrockmicro.com/) simultaneously with the clinical channels recorded with Micromed. In four HD participants, their HD ECoG data were recorded also via Micromed at 512 Hz (filtered at 0.15–134.4 Hz). In three of these patients, HD ECoG data were recorded simultaneously with and in addition to the clinical iEEG channels. In one patient, only HD or only clinical electrodes could be recorded at the same time, therefore no simultaneous data for HD and clinical iEEG is available for this patient. Instead, the patient performed the task twice: once which clinical iEEG data recorded, and another time with HD ECoG data recorded. The resting state data of this patient were also acquired asynchronously.

Additional behavioral recordings including electrocardiogram, electromyography, electrooculogram and respiration rate were collected as part of the clinical trajectory and are available for some patients.

#### FMRI data acquisition

Functional images were acquired on a Philips Achieva 3T MRI scanner using 3D-PRESTO^38^. Whole brain images were acquired with the following parameters: TR/TE = 22.5/33.2, time per volume 608 ms, FA = 10°, 40 slices, FOV = 224 × 256 × 160 mm and voxel size of 4 mm.

#### Structural data acquisition

For most participants the structural T1 images were acquired on a Philips Achieva 3T MRI scanner using TR/TE 8.4/3.2 ms, FA = 8°, 175 slices, FOV = 228 × 228 × 175 and voxel size of 1 × 1 × 1 mm. One participant had a 7T structural scan. Twenty participants had a 3T scan with different parameters from those described above: for example, sub-millimeter voxel size (thirteen patients), voxel size of 1 × 1 × 1.1 mm (three patients) or different number of slices (twenty patients).

### Data processing and curation

#### Localization of iEEG electrodes on structural images

Electrodes were detected on each patient’s post-operative computer tomography scan and coregistered to the anatomical MRI in native space. The electrode locations were corrected for brain shift and projected on a cortex surface rendering^39, 40^ using either SPM12 (Welcome Trust Centre for Neuroimaging, University College London, https://www.fil.ion.ucl.ac.uk/spm/) volumetric maps or Freesurfer (http://surfer.nmr.mgh.harvard.edu/) surface reconstructions.

We do not provide normalized electrode positions due to the difficulties in normalization and noticeable distortions in resulting positions. For visualization purposes only we projected individual electrode locations to Montreal Neurological Institute (MNI) space using SPM12 procedures of anatomical segmentation, normalization and image reslicing.

#### Identification of bad electrodes

A status (’good’ or ’bad’) is provided for each channel in the dataset. Channels with a ’bad’ status also have a description that explains why it was labeled ’bad’. Each channel was visually inspected with respect to signal outliers and artifacts. Noisy channels are marked ’noisy’ in the description of the channel status. Some electrodes were placed on top of another electrode grid or strip during implantation. These channels are marked ’on top’ and are not recommended for use in data analyses as they do not record directly from brain tissue.

#### Defacing of structural images

All structural images were defaced to comply with the requirements for sharing de-identified medical data.

### Data validation procedures

#### IEEG data validation

Data were preprocessed per subject using MNE-Python^41^ (https://mne.tools/). First, we selected channels of type “ECoG” and “sEEG”, and from them discarded previously identified bad channels. Then, a notch filter was applied to account for the line noise at 50 Hz and its harmonics. The data were then re-referenced using the common average signal, and band-specific neural signals were extracted using the Hilbert transform: delta (*δ*, 1 – 4 Hz), theta (*θ*, 5 – 8 Hz), alpha (*α*, 8 – 12 Hz), beta (*β*, 13 – 24 Hz) and the high-frequency band (HFB) component (60 – 120 Hz). Final envelopes were downsampled to 25 Hz.

As a basic check of the subjects’ response to the task we compared their neural activity during speech and music blocks. For this, we estimated an ordinary least squares fit to the HFB envelopes using the block design boxcar function. The fit and statistical analysis was performed using Python package statsmodels^42^ (https://www.statsmodels.org). Given a possible delay in the brain’s response to the auditory input and the possibility that this delay could vary across electrodes, we calculated the fit per electrode at all time lags within 1 second after the sound onset. The best fit across the lags was recorded together with the lag value. Significance of the fit was assessed parametrically based on the t-statistic for the block design regression weight. Here we report only positive t-statistics, which correspond to higher responses to speech and lower responses to music (for the block design predictor with zeros in music blocks and ones in speech blocks) that are significant at *p* < .01, Bonferroni corrected for the number of electrodes and lags.

In addition, per electrode we computed r-squared values (calculated as Pearson correlation coefficient squared and preserving sign of correlation) between speech and music blocks, speech blocks and task rest and speech blocks and natural rest. To reduce the number of multiple comparisons (number of electrodes × frequency bands) the analysis was only performed on the electrodes with a significant linear fit to the block design (see the analysis above). The three comparisons (speech vs music, speech vs task test and speech vs natural rest) were made separately for all extracted iEEG frequency bands. Significance of reported r-squared values was determined parametrically, reported values are significant at *p* < .05, Bonferroni corrected for the number of electrodes and frequency bands.

#### FMRI data validation

FMRI data were processed with FSL^43^ (https://fsl.fmrib.ox.ac.uk/). Preprocessing steps included estimation of motion parameters, detrending, low-pass filtering and spatial smoothing.

To assess basic data quality we first analyzed motion displacement plots (calculated using *fsl_motion_outliers* based on estimation of frame displacement^44^). We also computed the temporal signal-to-noise-ratio (tSNR) per voxel as the mean of the functional volumes over time divided by their standard deviation over time^45^. Nibabel^46^ (https://nipy.org/nibabel/) and Numpy^47^ (https://numpy.org/) Python libraries were used for this. To visualize average tSNR on the brain surface, we normalized individual subject’s tSNR maps and computed an average over all subjects per voxel in the MNI space. This map was projected on the standard average Freesurfer surface.

Then, subject-specific and group-level statistical analysis was performed to compare fMRI responses to speech and music. A general linear model was fitted to the fMRI data using the block design boxcar function and motion parameters as additional covariates. There analyses were performed using default parameters in FSL FEAT^48, 49^.

### Conversion to BIDS

We used in-house software to convert raw data files to the BIDS (fMRI) and iBIDS (iEEG) format. Code is available here: https://github.com/UMCU-RIBS/xelo2. Validation checks were performed using BIDS Validator (https://github.com/bids-standard/bids-validator), MNE BIDS routines (https://mne.tools/mne-bids/) and manual inspection of the iBIDS data.

## Data Records

The dataset is freely available at the openneuro.org database. All personal identifiable information has been removed and individual MRI scans have been defaced. The order in which subjects are presented in the dataset has been randomized and therefore does not follow any identifiable pattern (for example, alphabetical order or order by date of the experiment).

Data are organized according to the BIDS format^36, 50^. The root folder contains meta-information about the description of the dataset (*dataset_description.json);* the list of participants along with their demographic details, handedness and language dominant hemisphere (*participants.tsv*); the stimulus folder (*stimuli*) and individual data folders per participant (for example, *sub-01*, Figure 2a).

**Figure 2.**
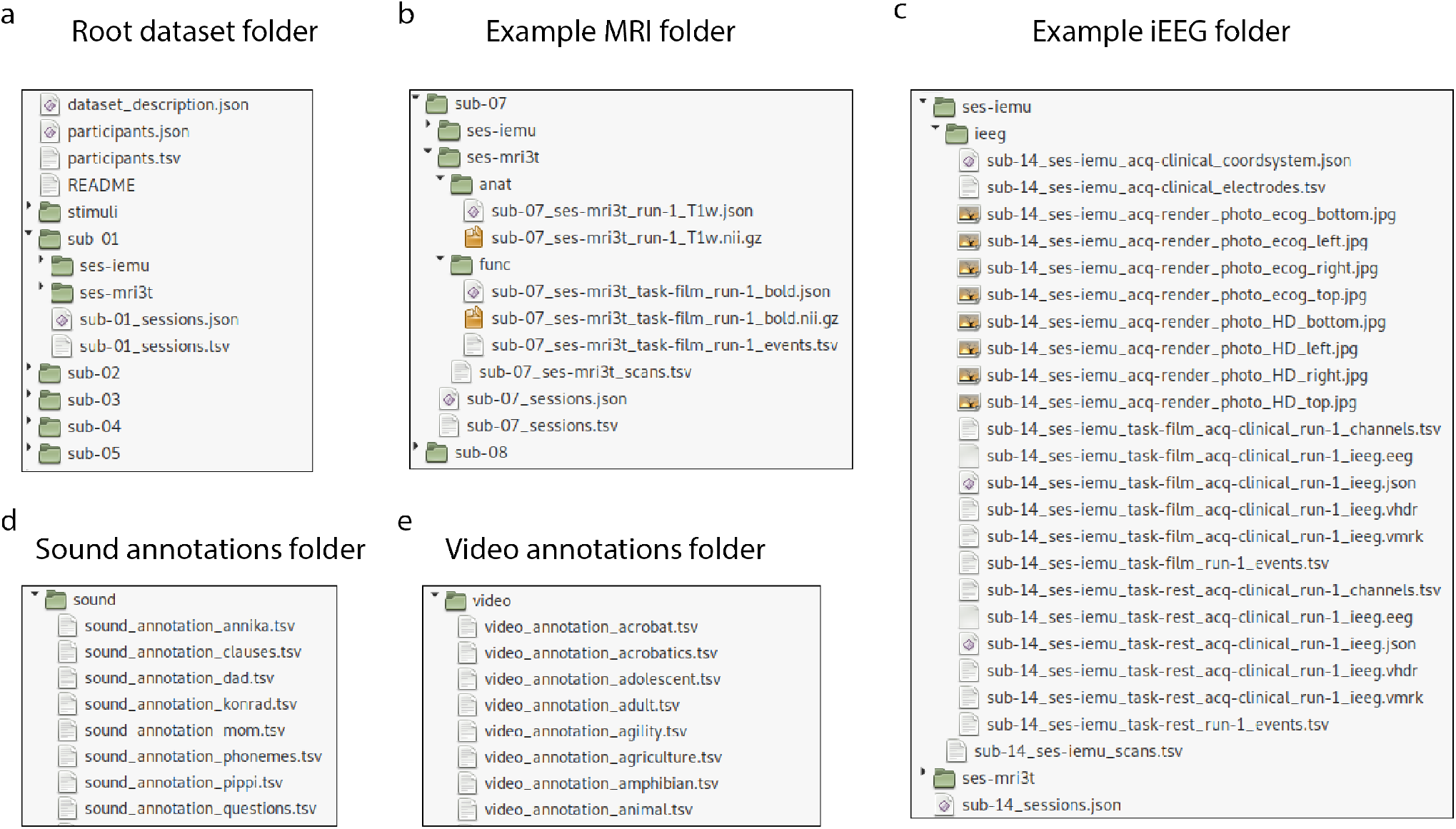
Overview of data records. a-d. Structure of folders and files in the dataset with example (f)MRI and iEEG folders from one subject.

### Stimulus folder

Two folders are provided under the Stimulus directory: *sound* and *video* (Figure 2d,e), each containing annotations for the corresponding stream of the film. The *sound* folder contains 13 tsv files. Each file contains soundtrack annotation with respect to the feature in the name of the file. For example, *sound_annotation_words.tsv* is the annotation of onsets and offsets for individual words. Each file has three columns: item (for example, individual words, phonemes, etc. depending on the feature), its onset and offset in seconds. There are seven annotations for language features: phonemes, syllables, words, verbs, clauses, sentences and questions; and six annotations for individual story characters: Pippi, Annika, Tommi, Mom, Dad and Konrad.

The *video* folder contains 135 tsv files: 129 for individual visual concepts and 6 for individual story characters. The list of visual concept was determined based on the visual concept recognition model, which automatically labelled each frame with visual objects and concepts it had been trained to detect (see Methods for more detail). Each file contains two columns: onsets and offsets in seconds.

### Participant data folders

Each participant’s folder contains one or two directories depending on what brain recordings were available. For patients who have both fMRI and iEEG data the folder contains two directories corresponding to (f)MRI and intracranial recording sessions respectively (for example, *sess-3t1* and *sess-iemu1*). For patients who only have iEEG data the folder still contains two directories, and the MRI directory only contains a structural MRI scan. For patients who only have fMRI data, there is only one directory corresponding to the (f)MRI session. Individual details of (f)MRI and intracranial data sessions can vary across participants. For example, one patient has a 7T MRI scan and therefore their (f)MRI folder is named *sess-7t1*.

#### IEEG folder

If available, iEEG recordings are stored in the patient-specific iEEG folder (for example, *sess-iemu1*). The folder contains all iEEG-related information including

- Locations of clinical iEEG (\**acq-clinical*\*) and HD ECoG (\**acq-HGgrid*\*) electrodes together with a sidecar json file that contains electrode metadata. Since both sEEG and clinical ECoG are acquired through the clinical setup, their electrode locations are stored together in the \**acq-clinical*\* file and can be differentiated by the column ’type’. In three HD participants their HD data were recorded through the clinical setup and therefore are part of the ’acq-clinical’ files. These HD ECoG electrodes can be identified based on the column ’size’ that represents the recording surface area (*mm*^2^), and is typically ~1 mm in HD electrodes. Electrode locations are provided in the native space and are the same for both movie-watching and resting state tasks.
- Rendering of electrode locations (\**rendering.tiff*) per acquisition type (’clinical’ and ’HDgrid’), in native subject space.
- Per task (’film’ and ’rest’) and when available, per acquisition type (’clinical’ and ’HDgrid’), a file with montage of the recording channels (\**channels.tsv*). In rare cases the montage differs between the tasks. The file contains information about iEEG electrodes and additional recorded channels (for example, marker channel, respiration rate, electrooculography, etc.). Per channel signal acquisition details are provided, including units of measurement, sampling frequency, channel status and others. Channel status indicates which electrodes are recommended for analyses (good) and which are not (bad).
- Per task (’film’ and ’rest’) and when available, per acquisition type (’clinical’ and ’HDgrid’), a file with experimental events (\**events.tsv*). For the movie-watching task (’film’) the file contains onsets and offsets for the task start, each music and speech block and the task end. For the resting state data (’rest’) the file contains onsets and offsets of the three-minute rest period that came either from the task logs (in the resting state task) or were manually annotated (in natural rest).
- Per task (’film’ and ’rest’) and when available, per acquisition type (’clinical’ and ’HDgrid’), main files with iEEG recordings in the BrainVision format (\**ieeg.eeg*, \**ieeg.vmrk*, \**ieeg.vhdr*).

#### (F)MRI folder

All participants have a folder that corresponds to the (f)MRI recording session (for example, *sess-3t1*). The folder contains an anatomical MRI scan (*anat* directory) and functional MRI data (*func* directory) from the movie-watching experiment if available. (F)MRI data are provided in the NIFTI format with sidecar json files that store additional metadata. Functional images are accompanied by the \**events.tsv* file that contains onsets and offsets of speech and music blocks of the movie-watching task measured in seconds.

## Technical Validation

### IEEG data validation

#### Bad iEEG channels

We made a summary of bad channels and channels recommended for analyses. Channels marked as bad are also excluded from the results presented below. These electrodes either were considered noisy based on the visual inspection of the data, or were located on top of other electrodes based on the photographs from the implantation or explantation surgeries. Only four participants had more than 10% of their intracranial electrodes marked as bad channels, whereas the median number of good channels across participants is 79 (Figure 3a).

**Figure 3.**
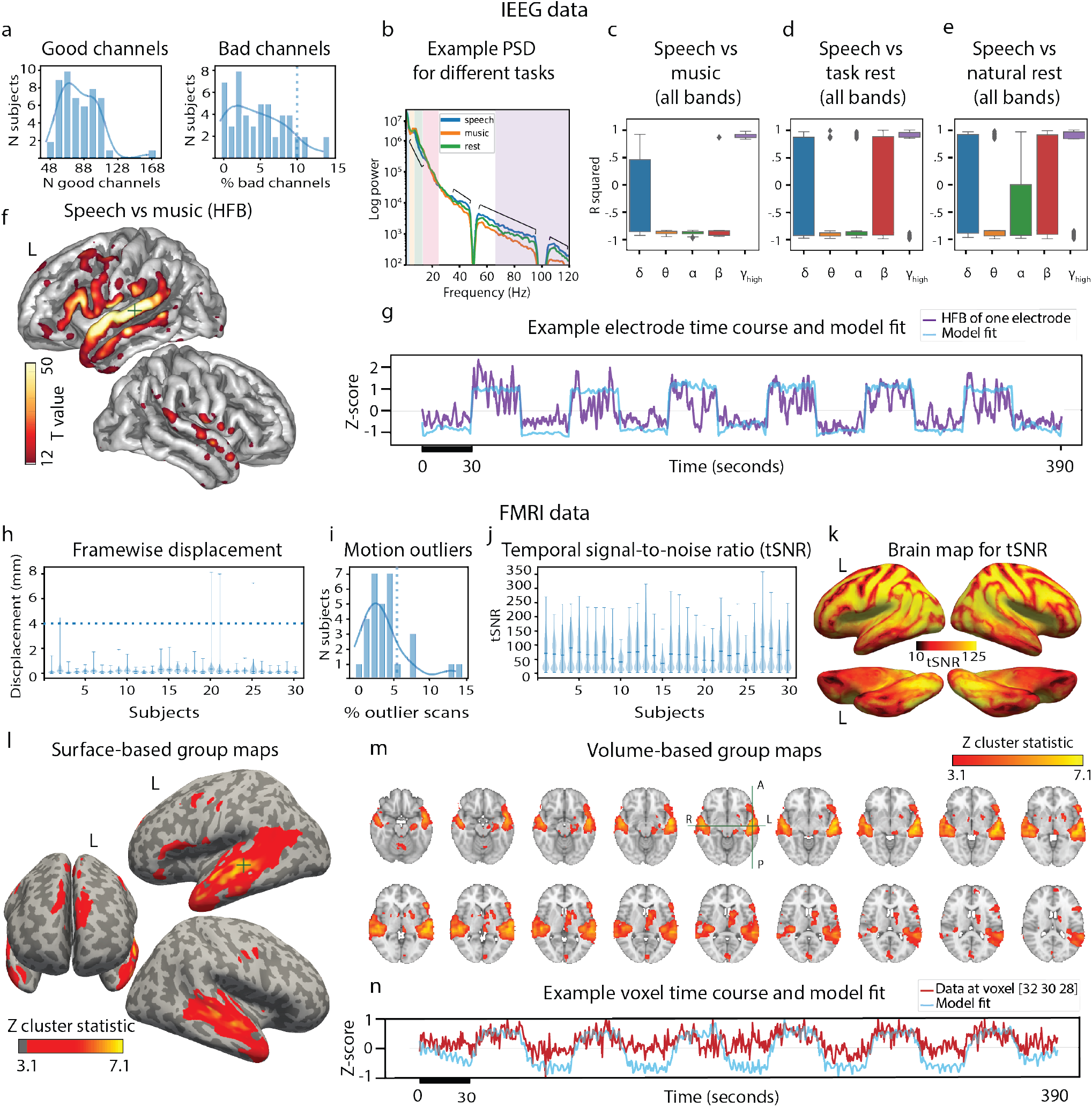
Technical data validation in iEEG (a-g) and fMRI (h-n). a. Histograms of good and bad channels over all subjects with estimated distribution density (solid line) and cut-off at 10% of bad channels (dotted line). b. Example of power spectral density (PSD) plot for one electrode in one subject per condition: speech, music and rest. Different frequency bands are highlighted (*δ*, *θ*, *α*, *β* and high frequency band, HFB). c-e. Boxplots of r-squared values (significant at *p* < .001) for three comparisons in the neural data: speech vs music (c), speech vs task rest (d) and speech vs natural rest (e), separately per frequency band. Boxes outline 25th and 75th quantiles, fliers show outliers (< 5% of data). f. Brain map (all patients) for results of the linear regression of HFB data to the block design in the movie-watching task (speech & music blocks). Only positive t-statistics significant at *p* < .001 are shown. For visualization a 2d Gaussian kernel (10 mm in width) was applied to each electrode’s central coordinate. Example time course fitted by the model (block design + audio envelope) and observed HFB in one electrode of one subject. h. Framewise displacement in fMRI data. Violin plots show entire data range, horizontal lines show medians. Dotted line shows voxel size (4 mm). i. Histogram of motion outliers with estimated distribution density (solid line) and cut-off at 5% of bad volumes (outliers determined by *fsl_motion_outliers*). j. Violin plots of the temporal signal-to-noise ratio (tSNR), same display setup as in h. k. Brain map of tSNR averaged over all subjects and projected onto the average surface. l. Group-level statistics based on results of the first-level analysis fitting a general linear model on fMRI data using the block design (with default FSL parameters). m. Same group-level brain map in volume slices. n. Example time course fitted by the model (block design + motion parameters) and observed fMRI in one voxel of one subject.

#### Response to the audiovisual movie task

Smaller subsets of the present dataset have previously been analyzed with respect to the auditory and visual processing of the movie stimulus^27, 51–53^. Here, we only show some basic results regarding the overall iEEG response to the task.

First, we performed a simple regression analysis that compares HFB responses during speech and music blocks. We mapped the resulting positive t-statistics that were significant at *p* < .001 (and subsequently Bonferroni-corrected for the total number of electrodes) onto the brain surface. This map showed preference to speech blocks over music throughout the perisylvian regions bilaterally and in inferior frontal gyrus, premotor and motor cortices on the left hemisphere (Figure 3f).

In addition, we computed mean power changes in frequency bands other than HFB and compared them across different conditions: speech, music and rest. We calculated r-squared values for three comparisons: speech vs music, speech vs task rest and speech vs natural rest, separately for delta, theta, alpha, beta and HFB mean power signal. The reported values are significant at *p* < .05 (Figure 3c-e). Consistent with the literature, the average pattern of HFB response was opposite to that of the lower frequency bands (theta and alpha) in all comparisons^54–56^. One notable difference between speech vs rest and speech vs music comparisons was stronger presence of positive r-squared values in the beta band during rest.

#### Resting state task and natural resting state data

To offer some form of comparison between rest data from a task and natural rest data from continuous 24/7 recordings, we report r-squared values for two comparisons: speech vs task rest and speech vs natural rest (Figure 3d-e). Both plots look very similar across almost all frequency bands (except for alpha) and suggest that either rest can be used as a baseline or control condition for investigating speech responses. Further investigation of the similarities and differences between the two sources of resting state data is an interesting research venue that the present dataset easily lends itself to.

### FMRI data validation

#### Analysis of motion

Based on the motion parameters obtained as part of the FSL preprocessing pipeline we calculated framewise displacement^44^ of each participant’s head in the scanner (Figure 3h). The analysis showed that overall there was little motion above one voxel size. Analysis of outliers based on motion showed that only five participants had more than 5% of their functional volumes marked as outliers (Figure 3i). The amount of motion in these five patients may be considered excessive compared to healthy volunteers, however it is common in the clinical population^57^. The five participants with somewhat excessive motion had iEEG data from the same experiment and were therefore included in the dataset. Many methods for excessive motion correction exist and we refer readers to some of them in Usage Notes.

#### Temporal signal-to-noise ratio

Temporal signal-to-noise ratio is a measure of signal dropout and effects of noise over time. It can be used to estimate how much scanning time is necessary to detect statistical effects of varying strength in the data^45^. It is known that block designs provide a robust method for observing reliable activation patterns, but the movie stimulus also contained more sparse auditory and visual events. Given high enough tSNR, fMRI data can be analysed with respect to such individual events. We calculated that the mean whole-brain tSNR across participants was 66.34 ± 16.13, not much lower than the value typically reported in fMRI datasets (≈ 70) with healthy volunteers and a smaller voxel size^7, 58^ (Figure 3j). Inspection of the brain maps revealed a typical pattern with lower tSNR values for the anterior temporal lobe and the orbitofrontal cortex (Figure 3k). Lower tSNR values were also observed dorsally around the sulci. This could be due to use of the PRESTO scan sequence.

#### Response to the audiovisual movie task

We also performed a simple analysis to estimate overall participants’ response to the audiovisual movie task. For this, we fitted a general linear model per participant and estimated the group effects for the contrast comparing speech and music blocks. Group statistic shows a strong effect in brain areas typically associated with auditory and language processing including bilateral superior temporal gyrus, left inferior frontal gyrus, bilateral precentral gyrus and bilateral supplementary motor cortex (Figure 3l-n).

### Usage Notes

The dataset can be downloaded from the open public repository at https://openneuro.org/datasets/ds003688. Under the Public Domain Dedication and License, the data are freely available for non-commercial research purposes. Below we summarize a number of things to keep in mind when working with this dataset.

### IEEG data

#### Known issues

- The iEEG electrode coverage is much denser in the left than the right hemisphere. This should be taken into account when interpreting results of future analyses. In some cases it may be a good idea to confine iEEG analyses to the language-dominant hemisphere. Combining iEEG with whole-brain fMRI data may be useful when addressing inter-hemispheric differences.
- The present dataset includes a few difficult cases where accurate estimation of electrode locations was tricky. A small number of patients had an earlier (i.e. before the iEEG implantation) tissue resection followed by a build-up of liquid in the resection cavity, or a pathology (for example, a tumor) that may have affected the tissue under the electrodes. In addition, in patients who only had sEEG electrodes (three patients), their CT scan (used for electrode localization) lacked resolution for accurate separation of center of mass for every electrode.
- In one participant (’sub-44’) it was not possible to record HD and clinical ECoG simultaneously as both types of electrodes were recorded through the Micromed system that is limited to 128 channels. Therefore, the montage needed to be changed to switch between the recordings. Movie-watching and rest data in this participant were recorded twice: once with clinical ECoG and another time with HD ECoG grid.
- In one participant (’sub-29’) there was no temporal synchronization in the resting state recordings between HD and clinical ECoG data. This was due to an error in the recording setup.
- One participant (’sub-32’) is missing resting state iEEG data. There were no 24/7 continuous recordings of this patient available from the clinic.

#### Additional notes

- It has been shown that, in general, iEEG responses recorded from patients with epilepsy reflect states similar to healthy controls^59^, yet it is possible that some individual patient’s data can be affected by epileptic or interictal events.
- In preparation of this dataset we identified bad channels in each participant’s recordings. This was done based on visual inspection, calculation of basic statistics of the signal (mean signal and its variance) and photographs from implantation or explantation surgeries. Several alternative methods have been proposed to automate the process, and we encourage users to explore them^60, 61^.
- HD ECoG and sEEG data are advised to be processed separately from clinical ECoG data. For sEEG, bipolar reference and taking into account grey and white electrodes may be preferred^62, 63^. HD ECoG data allows to zoom in on the neural processing in one specific region (typically, sensorimotor cortex), and it is therefore best to use either local or separate common average reference for it. These recordings are also often of higher sampling rate (2000 Hz) and this can be leveraged for HFB analyses.
- Physiological measures among the iEEG channels (electrocardiogram, breathing and electrooculography), if available, are a valuable source of information. Previously we used electrooculography to infer saccade data, although patterns of eye blinks have been related to processing of relevant detail in perceptual input^64^. Electrocardiogram and breathing have been shown to correlate with cognitive states during experimental stimulation^65–67^.
- Resting state data is another useful source of information. It can be used as a baseline for the task data and it can also be studied on its own by exploration of the internal dynamics in the task-free neural activity.

### FMRI data

- PRESTO scans have superior temporal resolution compared to the standard EPI sequence^38^. Some caution is advised in processing and interpreting PRESTO data. For example, there has been conflicting evidence about inclusion of motion parameters in modeling fMRI data based on experimental variables (for example, using a general linear model).
- In four fMRI participants amount of estimated motion exceeded one voxel size (4 mm). This excessive motion is due to the fact that all fMRI data comes from epilepsy patients. This is intentional as fMRI data is meant to be complementary to the iEEG recordings, and here we provide data from a considerable number of patients who did the same task with both recording modalities. However, it is known patients are less likely to remain stationary in the scanner to the same degree healthy volunteers would. Several methods to account for this motion have been proposed and successfully used to mitigate the issue, including motion scrubbing^68^, Volterra expansion for general linear models^57, 69^, independent component analysis for artifact removal^70^ and other despiking and denoising methods. We can recommend that users explore software that incorporates advanced motion correction methods such as fMRIprep^71^ and ArtRepair^72^ toolbox for SPM.

### Other issues

- The soundtrack of the audiovisual film was originally in Swedish. For all our experiments with Dutch patients, many of whom were children, we used a movie version dubbed into Dutch.
- Four participants (’sub-11’, ’sub-37’, ’sub-49’ and ’sub-63’) have missing information about their handedness.
- Four participants (’sub-01’, ’sub-11’, ’sub-30’ and ’sub-33’) have missing information about their language dominance hemisphere.

## Code availability

The code used to perform technical validation on the iBIDS dataset is available at https://github.com/UMCU-RIBS/ieeg-fmri-dataset-validation. We also provide a set of utility scripts to get the new users started with processing and visualizing the data (https://github.com/UMCU-RIBS/ieeg-fmri-dataset-quickstart).

## Acknowledgements

This work was supported by the European Research Council (Advanced iConnect Project Grant ADV 320708) and the Netherlands Organisation for Scientific Research (Language in Interaction Project Gravitation Grant 024.001.006). We thank Frans Leijten, Cyrille Ferrier, Geert-Jan Huiskamp, Sandra van der Salm and Tineke Gebbink for help with collecting data; Peter Gosselaar and Peter van Rijen for implanting the electrodes; the technicians and staff of the clinical neurophysiology department and the patients for their time and effort; and the members of the UMC Utrecht ECoG research team for data collection. We also thank the Swedish Film Institute film company for their help and the provided materials.

## Author contributions statement

All neural data were collected over a span of 14 years (2007 to 2021) at the University Medical Center Utrecht. NFR is the founder of the research lab. MJV, EA and NFR started clinical screening and research with iEEG patients in 2007. All other authors became involved in data collection, curation and organization at later times as they joined the lab. In 2019 JB and NFR conceived the idea of sharing the accumulated data with the scientific community. MJV, MPB and JB worked on obtaining patients’ permissions to share their anonymized data. MPB developed electrode localization and coregistration techniques and obtained electrode coordinates for many patients. JB and MJV collected all metadata. JB and ZVF assembled and curated the data. GP converted the data into (i)BIDS. JB performed technical data validation and (i)BIDS conversion checks, and wrote the first draft of the paper. All authors participated in the discussion of conceptual and technical details, and contributed to writing and editing of the manuscript.

## Competing interests

The authors declare no competing interests.

## Notes

https://openneuro.org/datasets/ds003688

### Competing Interest Statement

The authors have declared no competing interest.

